# MB-GAN: Microbiome Simulation via Generative Adversarial Network

**DOI:** 10.1101/863977

**Authors:** Ruichen Rong, Shuang Jiang, Lin Xu, Guanghua Xiao, Yang Xie, Dajiang J. Liu, Qiwei Li, Xiaowei Zhan

**Affiliations:** Quantitative Biomedical Research Center, Department of Population and Data Sciences, University of Texas Southwestern Medical Center, Dallas, Texas, 75390, USA; Department of Statistical Sciences, Southern Methodist University, Dallas, TX 75275, USA; Harold C. Simmons Cancer Center, University of Texas Southwestern Medical Center, Dallas, Texas, 75390, USA; Department of Bioinformatics, University of Texas Southwestern Medical Center, Dallas, Texas, 75390, USA; Institute for Personalized Medicine, College of Medicine, Pennsylvania State University, Hershey, Pennsylvania, 17033, USA; Division of Biostatistics, Department of Public Health Sciences, College of Medicine, Pennsylvania State University, Hershey, Pennsylvania, 17033, USA; Department of Mathematical Sciences, The University of Texas at Dallas, Dallas, Texas, 75080, USA; Center for the Genetics of Host Defense, University of Texas Southwestern Medical Center, Dallas, Texas, 75390, USA

**Keywords:** Microbiome simulation, Generative adversarial network, Deep learning

## Abstract

Simulation is a critical component of experimental design and evaluation of analysis methods in microbiome association studies. However, statistically modeling the microbiome data is challenging since that the complex structure in the real data is difficult to be fully represented by statistical models. To address this challenge, we designed a novel simulation framework for microbiome data using a generative adversarial network (GAN), called MB-GAN, by utilizing methodology advancements from the deep learning community. MB-GAN can automatically learn from a given dataset and compute simulated datasets that are indistinguishable from it. When MB-GAN was applied to a case-control microbiome study of 396 samples, we demonstrated that the simulated data and the original data had similar first-order and second-order properties, including sparsity, diversities, and taxa-taxa correlations. These advantages are suitable for further microbiome methodology development where high fidelity microbiome data are needed.

## 1. Introduction

Microbiome is a collection of trillions of microorganisms living within humans. Previous studies have revealed that the microbiome has a profound impact on human disease, including inflammatory bowel disease, colorectal cancer, type-2 diabetes, and psychiatric disorders^[1,2,3,4]^. A powerful and increasingly popular method to study the microbiome and disease is metagenome-wide association study (MWAS). It utilizes the taxonomical abundance data of thousands of bacteria generated from sequencing instruments, and calculates the association strengths between the bacterial abundances and the phenotypes. As researchers have expanded their interests into studying microbial associations in human physiology, MWAS has taken on an increasingly critical role.

A successful MWAS relies on valid statistical models^[5]^, and the evaluation of MWAS models relies on simulations. As microbiome data have unique characteristics, it is not trivial to simulate microbiome abundances with high fidelity to the real data. For example, microbiome abundances are sparse, over-dispersed (large variances compared to means), and have an intrinsic phylogenetic relationship^[6]^. In addition, as the microbiome consists of interactive communities, the microbiota form complex taxa-taxa relationships with a noneligible second-order covariation^[7,8]^. However, a simulation method that can capture the first order (e.g., sample-level) characteristics while maintaining the second order (e.g., taxa-taxa level) relationships is lacking in the current literature. For example, explicit statistical distributional assumptions, such as the zero-inflated logistic normal distribution, were introduced to simulate individual bacterial taxon^[9]^. Although the simulated data have the desired sample-level and taxa-level properties (e.g., sparsity and overdispersion), they ignore the taxa-taxa relationships. Other methods, such as Normal To Anything (NorTA)^[10]^, have attempted to model the taxa-taxa relationships, but its performance at the sample-level is not satisfactory, as we will demonstrate in this manuscript.

Given the above challenges in explicitly modeling the microbiome abundances, we seek a novel deep learning-based approach to implicitly compute simulated microbial abundances with desired sample-level and taxa-taxa interactive characteristics. We refer to this simulation framework as MBGAN (Microbiome Generative Adversarial Network), as it is adapted from a Generative Adversarial Network (GAN) framework^[11]^. The foundational work on GANs was proposed in 2014^[11]^. GANs have a generator network and a discriminator network. The generator network takes random noise and outputs simulated data. The discriminator network takes both the simulated and the real datasets, and classifies the inputs as real or fake. In the network training stage, the generator focuses on increasing the similarity between the simulated data and the real data; meanwhile, the discriminator works on better distinguishing between the simulated and the real data. When the training stage finishes, the generator will be able to simulate data that are hard to be distinguished from the real data by the discriminator. Due to the excellent performance of GANs compared with conventional approaches (e.g., variational auto-encoder^[12]^), GANs have wide applications such as image synthetization (e.g., human facial images^[13]^), text generation (e.g., visual paragraph generation^[14]^) and music synthetization (e.g., music composition^[15]^). Recently, GANs have also been adapted for biomedical research. For example, GANs have been applied in generating sequence data (e.g., t-cell receptor sequences^[16]^) and enhancing medical imaging^[17]^. Since GANs have achieved impressive success in simulation tasks, we are motivated to adapt them to simulate high fidelity microbiome datasets.

In this manuscript, our contribution in MB-GAN is the adaption of the GAN framework for simulating the microbiome data. Compared to the existing GAN framework, our algorithm can reach convergence efficiently and is easily applicable, as it can simulate new datasets based on a set of input microbiome data without explicit modeling. We demonstrated that the simulated datasets have similar microbiome data characteristics, including both of the first order (sample-level properties such as sparsity and diversity) and the second order properties (taxa-taxa correlations). The simulated data can be used in further methodology development, such as for MWAS, by imposing proper disease models.

## 2 Methods

### 2.1 Construction of MB-GAN

We created MB-GAN based on the Wasserstein GAN with gradient penalty (WGAN-GP) framework^[18]^, as it exhibited several improvements over the previous work. Briefly speaking, traditional GANs suffer from several practical problems due to the design of its binary discriminator and loss functions. Two major concerns are: 1) the generator will stop updating when discriminator is overpowered (e.g., generator’s gradients vanish or explode); 2) generated samples lack enough variability (model collapse^[19]^). These problems make traditional GANs hard to train and thus the powerful generative framework of GANs had limited usage. WGAN^[20]^ modifies the original binary discriminator into a critic that measures how good the data are by a scalar similarity score (e.g., the Earth Mover’s distance). Comparatively, the discriminator measures whether the data are real or fake by probabilities. By taking the advantages of Earth Mover’s (EM) distance, WGAN provides a smoother gradient everywhere and thus keeps the generator learning new knowledge even when the critic already performs well. WGAN-GP^[18]^ further reforms the gradient clipping into a gradient penalty to solve a practical issue lead by WGAN’s Lipschitz constraint. So far, WGAN-GP has no sign of any gradient problems or model collapse and thus is an optimized framework for training a well-performed GAN generator.

The detailed architecture of our proposed MB-GAN network structure is illustrated in Figure 1. It is built on top of WGAN-GP with an additional weighted phylogenetic transformation layer. In the training phase, the network requires a real microbiome dataset *x*, which contains the relative abundances of taxa observed across different samples. For each sample, the relative abundance of a taxon is defined as its percent composition relative to the total abundances observed in that sample. The generator will take random noise inputs (e.g., *z ∼ N* (0, 1) as Gaussian noise) and output simulated microbiome datasets (*g_θ_*(*z*)). Both real microbiome samples and simulated microbiome samples will be combined in one batch. Then all data will undergo weighted phylogeny transformation, and be sent to a critic to calculate the EM Distance (*β*-diversity) between real and simulated samples. The generator and critic will be differentiated against the distance alternatively to update the model weights.

To handle microbiome data, we add a transformation layer. It integrates additional phylogenetic information into the critic in order to biologically measure the dissimilarity between the real sample and generated sample. Specifically, the transformation layer expands the species level table based on the phylogenetic tree to make a full abundance matrix for all taxa from phylum to species. When feeding the transformed matrix into WGAN-GP framework, the critic calculates the EM distances along the given phylogenetic tree. In the microbiome literatures, these values are equivalent to non-normalized Unifrac *β*-diversity^[21]^. Thus our MB-GAN framework utilizes the biologically meaningful Unifrac *β*-diversity between the true and simulated microbial environment to score the quality of the generative model. In addition, taxa weights such as tree branch lengths and normalization methods such as logarithmic transformation can be integrated into the transformation layer as a generalized-weighted-Unifrac critic^[22]^. For example, we applied logarithmic transformation as the normalization to our MB-GAN framework. In practice, we observed that the log transformation provided robust and satisfactory results regarding the first and second order properties, even without the help of tree branch length.

**Figure 1:**
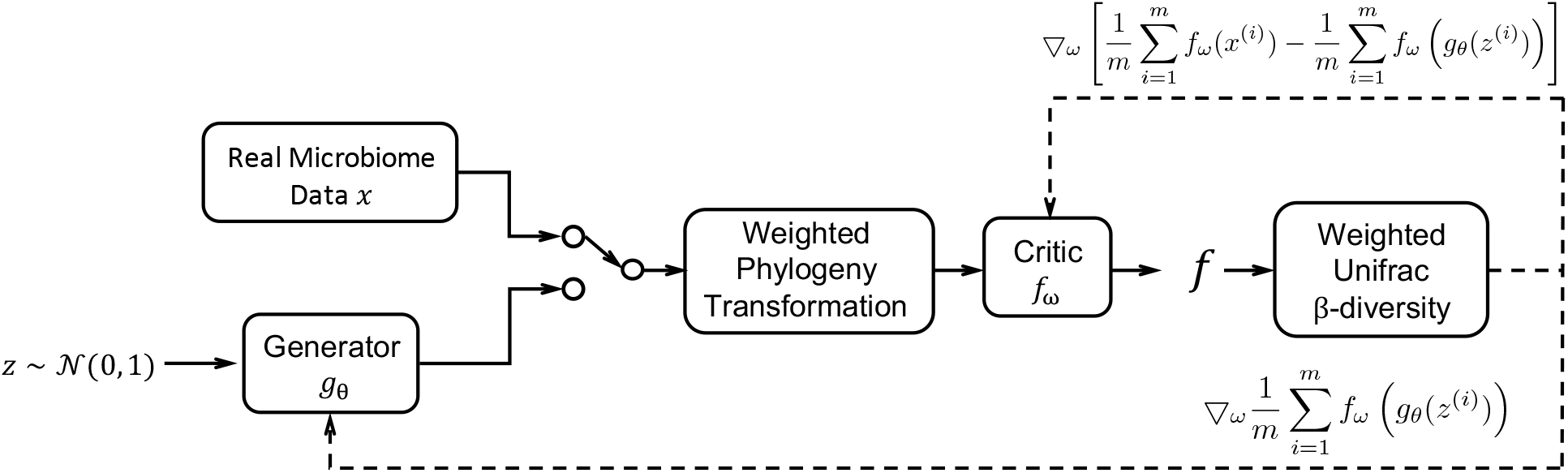
The MB-GAN network architecture. For the solid lines: the Gaussian noise *z* is passed to the generator model *g_θ_*; both real microbiome data *x* and simulated microbiome data *g_θ_*(*z*) undergo weighted phylogeny transformation and are passed to the critic model *f_ω_*; the function *f* calculates the weighted Unifrac *β* diversity score. For the dashed lines: the differentiated scores will be used to update generator weights *θ* and critic weights *ω* at the *i*-th iteration.

### 2.2 Implementation

#### 2.2.1 Data processing

As illustrated in Figure 1 and Supplementary Figure 1, the input of the generator model is a Gaussian random noise vector, and the inputs of the critic model are the real data and the simulated data. Next, both the real and the simulated data will go through the weighted phylogeny transformation, which utilizes a rooted taxonomic tree (or phylogenetic tree) to expand the species (or OTUs) level abundance matrices. The outputs include the abundance of all internal nodes for each input sample. Last, the critic uses the outputs to calculate a Unifrac diversity score. The score is essentially the EM distance along the given taxonomic tree, and can represent the performance of the generator model.

#### 2.2.2 MB-GAN training

We incorporate the training method in WGAN-GP with our own customization. For the critic, we use Wasserstein loss to measure the EM distance between the real data and the simulated data. Meanwhile, we employ the GP loss to achieve Lipschitz constraint required by WGAN-GP. We weight the Wasserstein loss and GP loss by 1:10. For the generator, we use Wasserstein loss to measure the minimum cost to change the simulated data into real data. In the MB-GAN training phase, we use RMSprop optimizer with learning rate 5 *×* 10^*−*5^ to update the model weights. The generator and critic will be differentiated alternatively, with 5 steps of critic followed by 1 step of generator in each iteration.

We assess the convergence of MB-GAN using two statistics. The first statistic combines the Wasserstein loss of generator and the Wasserstein loss of critic. Indeed, these losses can also be used for model selection. The second statistic is the mean of the mean square error (MSE) of the change of model parameters across iterations. The training of critic and generator is considered to reach convergence when the difference of the mean MSE is less than 1 *×* 10^*−*8^ in a 1000-iteration window. In practice, it is not necessary to stop the model training when the mean MSE is small. As WGAN-GP shows no sign of collapse during the training, it is recommended to train the model as long as possible even when the model reaches convergence, and then the best model can be selected by comparing model performance.

### 2.3 Comparison and evaluation of MB-GAN

#### 2.3.1 Comparison with Normal To Anything

We compare MB-GAN to Normal To Anything (NorTA), an alternative simulation methods for microbiome data. NorTA can generate multivariate random variables with a pre-specified correlation structure^[10]^. In general, NorTA transforms a multivariate Gaussian random variable with a given correlation structure to an arbitrary discrete or continues random variable, where the transformation is determined by the marginal distribution of the target random variable. In recent microbiome network studies, NorTA has been used to generate high-dimensional sparse count data with the underlying correlation structure defined by a target correlation matrix^[23,24]^. Choosing the zero-inflated negative binomial (ziNB) as the target marginal distribution in the transformation step was shown to well characterize the overdispersion and zero-inflation observed in real microbiome data ^[23]^. Briefly, the NorTA method includes the following steps: 1) remove the taxa with zeros across all real data samples; 2) generate an *n × p* multivariate normal random variable with zero mean and *p × p* correlation matrix calculated from the real data, where *n* and *p* are the sample size and taxa number, respectively; 3) for each taxon *j*, apply a standard normal cumulative distribuion function transformation to get a uniform random variable; 4) apply the quantile function of a zero-inflated negative binomial (ziNB) distribution to each uniform random variable to generate the count vector for each taxon *j*, with the parameters of the ziNB distribution estimated from the observed count data of taxon *j*; 5) compositionalize the resulted *n × p* count matrix by dividing each sample (row) by the total counts in that sample. The above steps are implemented by the R package SpiecEasi^[23]^.

#### 2.3.2 Evaluation of model performance

To evaluate the quality of the simulated datasets, we compare sample-level statistics and taxa-taxa correlations calculated from the real data and the two types of simulated data.

We use sparsity and diversity to measure the sample-level characteristics. The sparsity is defined as the proportion of zeros in a sample. In MB-GAN outputs, abundances less than 1 *×* 10^*−*4^ will be truncated to zero. The *α*-diversity for each sample is defined as the Shannon index: 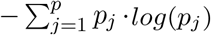. Here, we assume we have *p* taxa in total, and *p*_*j*_ is the relative abundance of taxon *j* in one sample. Then the *α*-diversities of the real and the simulated samples are compared through Wilcoxon rank-sum test. We used the *β*-diversity between two samples *m* and *k*, defined as their Bray-Curtis dissimilarity: 1 *−* 2 *× c_m,k_/*(*s*_*m*_ + *s*_*k*_). Here, *c_m,k_* is the sum of the lesser values for only those taxa in common between both samples, and *s_m_, s_k_* are the total number of taxa in sample *m* and *k*, respectively. We intentionally avoid using Unifrac *β*-diversity as it has already minimized by the MB-GAN model. Then the *β*-diversities are tested with the analysis of similarities (ANOSIM) and are visualized with the non-metric multidimensional scaling analysis (nMDS).

We use Pearson correlation coefficients between taxa to measure the taxa-taxa relationships. We exclude taxa with an excessive number of zeros (more than 90%) across all real samples. Then the pairwise correlation coefficients are calculated among all the remaining taxa.

## 3 Results

### 3.1 Simulate microbiome abundances from a real colorectal cancer study

We evaluate MB-GAN using a real sequencing data from a human gut microbiome study^[25]^. The data contains 396 sequenced shotgun metagenomic samples with 148 inflammatory bowel disease (IBD) patients and 248 healthy controls. The original sequencing data from the fecal samples are available in the European Bioinformatics Institute database with accession code ERP002061. We used curatedMetagenomicData^[26]^ to obtain the taxonomic abundance table of all samples with 1,939 detected taxa at different taxonomic levels.

To simulate microbiome data using MB-GAN, we used the 148 cases (or 248 healthy controls) as real data to train the MB-GAN network for 100,000 iterations (batch size set to 32). The generators and critics reached convergence in 10 minutes after 20,000 iterations (Figure 2). It is noteworthy that the traditional GAN failed to converge and the WGAN alone failed to simulate microbiome data with high variation. In contrast, MB-GAN converged rapidly in practice and provided high fidelity data for both case and control samples, as will be demonstrated next.

### 3.2 Evaluation on sample-level properties

To understand the simulation performance of MB-GAN, we first compare the sample-level statistics calculated from the real data and the simulated data (by MB-GAN or NorTA). The sample-level properties include sparsity, *α*-diversity and *β*-diversity. As NorTA dropped the taxa with zeros across all samples in its simulation process, the comparisons focus on the species shared among the real data and the two simulators.

First, we evaluate sample sparsity which is the proportion of zeros in a sample. For the real data, the observed sparsity ranges from 0.71 to 0.90, with median values being 0.80 and 0.83 for the case and the control group, respectively. As for the two types of simulated data, the lower bound of sample sparsity by MB-GAN matches well with the real data for both groups, but in general the MB-GAN simulated data show a slightly higher sparsity with a median of 0.83 and 0.85 for the case and the control group. NorTA simulated data, on the other hand, tends to underestimate the sparsity in both case and control groups. The median values of the sample sparsity are all below 0.8. for the two groups, and the maximum sparsity is less than 0.85. Thus MB-GAN simulated data better capture the sparsity observed across all the samples of the real data.

Next, we evaluate the *α*-diversities of the simulated data from MB-GAN or NorTA. We compared the Shannon indices calculated from the simulated and the real samples. The Shannon index is a metric that weights the relative abundance of species by their relative evenness in a sample. As an *α*-diversity index, it provides more information than simply species richness (i.e., the number of species in the sample) since it considers the relative abundances of different species. Therefore, a better matching between the Shannon indices calculated from the simulated data and the real data suggests that the simulator can better characterize the biodiversity of the real data. As shown by the box plots in Figure 3 (a) and (b), the Shannon indices calculated from MB-GAN simulation data match consistently with the real data in both the case and the control groups. The p-value given by the Wilcoxon rank-sum test is 0.83 for the case group and 0.57 for the control group. However, the Shannon indices by NorTA are clearly larger than the real ones, and the variation among the results is not well-characterized. This suggests that the samples simulated by NorTA are less diverse compared to the real data. Again, the Wilcoxon rank-sum test yields p-values less than 0.0001 for both groups when comparing the real data and NorTA simulated data. In all, our results suggest that the MB-GAN simulated data can better mimic the real microbiome data with respect to *α*-diversity.

**Figure 2:**
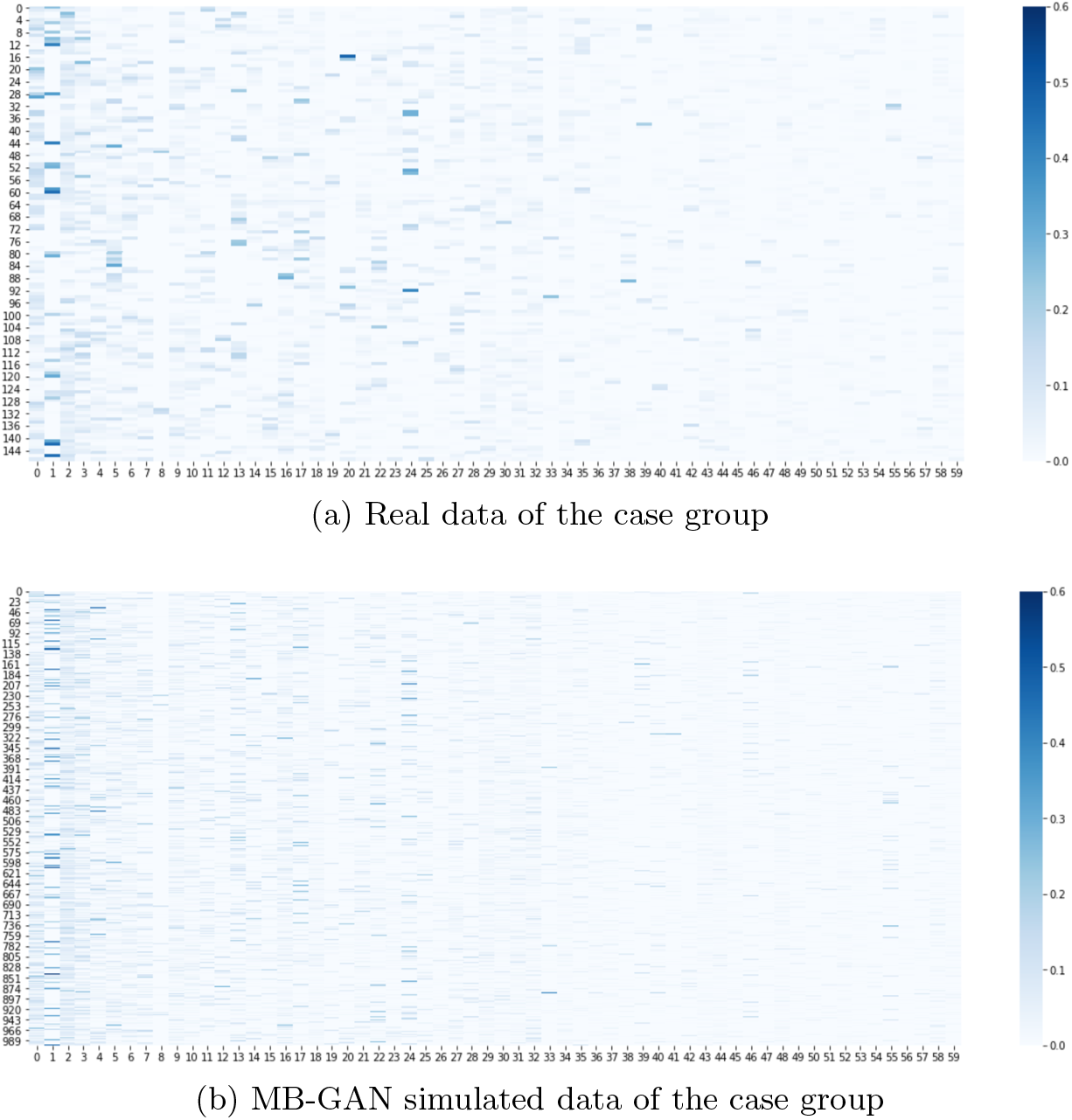
Heatmap of the 60 most abundant taxa from (a) the 148 samples in the case group of the real data, and (b) the 1,000 simulated MB-GAN case samples after 20,000 iterations

Finally, we evaluate the *β*-diversity of the simulated data from MB-GAN or NorTA, and compared them with the *β*-diversity of the real data. *β*-diversity measures how different samples are from each other, and a commonly used way to visualize *β*-diversity is by non-metric multidimensional scaling (nMDS) analysis. Here, since the samples are compared based on their species level abundances with no additional taxonomy information, we employed the Bray-Curtis metric to calculate the two-dimension nMDS values. Figure 4 visualizes the result from the case (left) and the control (right) group, respectively. In both scenarios, the clear overlap between the real data and the MB-GAN simulated data demonstrates the similarity between those samples. The NorTA simulated data, however, shows a unique circular pattern that is different from the real data or the MB-GAN simulated data. Furthermore, we test the similarity between samples from the real and the simulated data through analysis of similarities (ANOSIM)^[27]^, which is suitable for our situation since the real data and the simulated data have unbalanced sample sizes. In particular, we first calculated the distance matrices of the real and the simulated data using the Bray-Curtis metric. Then, we performed ANOSIM with 1,000 permutations. The results are summarized in Table 1. A small p-value of 0.001 suggests significant difference between the samples generated by NorTA and the real data, for both the case and the control groups. The MB-GAN simulated data, on the other hand, shares more similarity with the real data in both groups.

**Figure 3:**
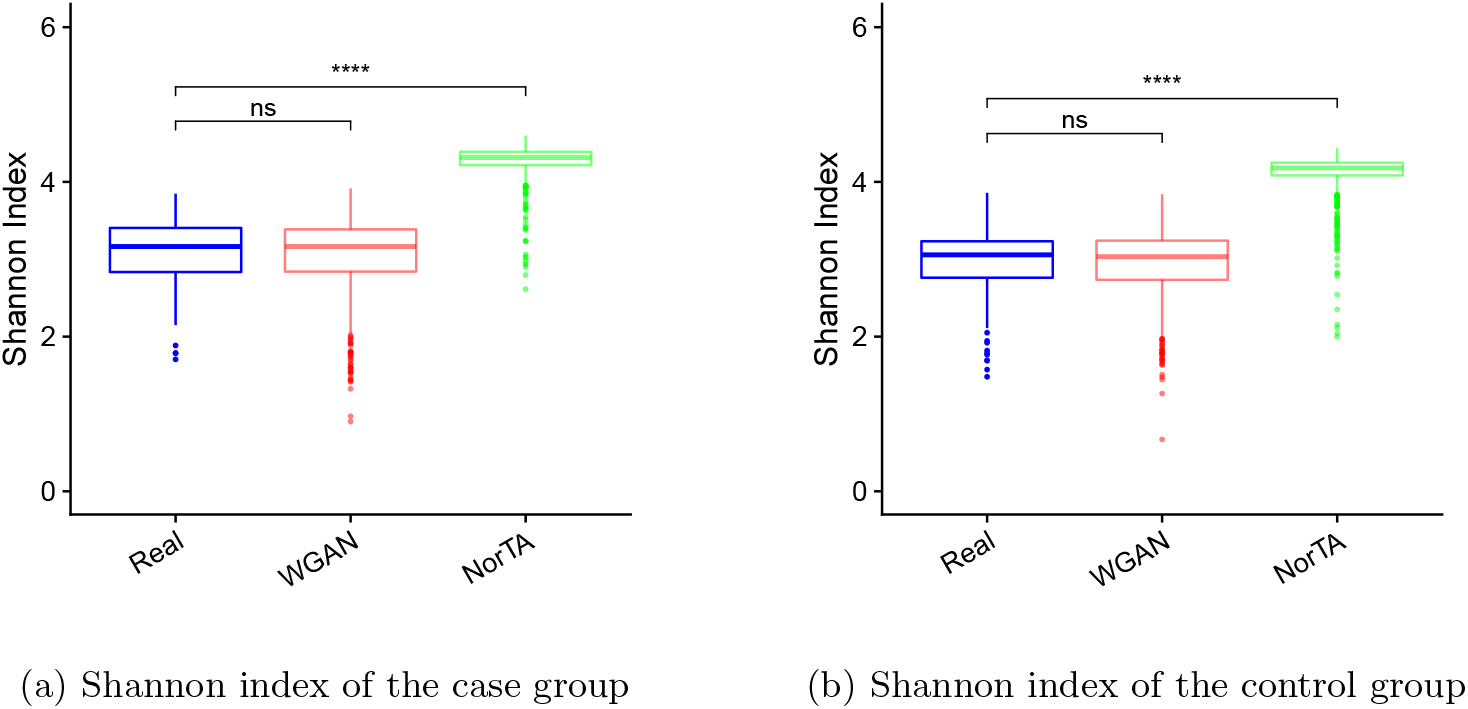
Box plots of Shannon index calculated from three datasets (Real, MB-GAN and NorTA) under (a) case or (b) control group.

**Table 1:**
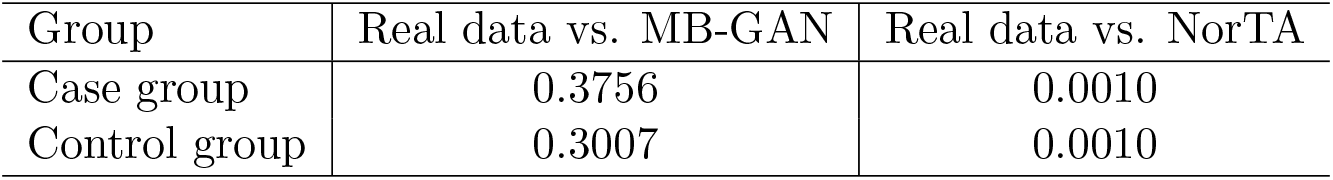
P-values from analysis of similarities (ANOSIM) between the real data and the simulated data from MB-GAN or Normal To Anything (NorTA). The null hypothesis is that there is no significant difference between two groups of samples.

| Group | Real data vs. MB-GAN | Real data vs. NorTA |
| --- | --- | --- |
| Case group | 0.3756 | 0.0010 |
| Control group | 0.3007 | 0.0010 |

### 3.3 Evaluation on taxa-taxa relationships

In addition to comparing the sample-level similarities, we illustrate that MB-GAN has the advantage of preserving the second order characteristics in the real data, as measured by the correlation structure among the taxa. We visualize the correlation matrices and the empirical distributions of the correlation coefficients to compare the real data and the two types of simulated data (MB-GAN and NorTA). Here, we only use the species-level compositional data with less than 10% zeros across all samples, since these taxa contain more information in capturing the taxa-taxa interactions.

**Figure 4:**
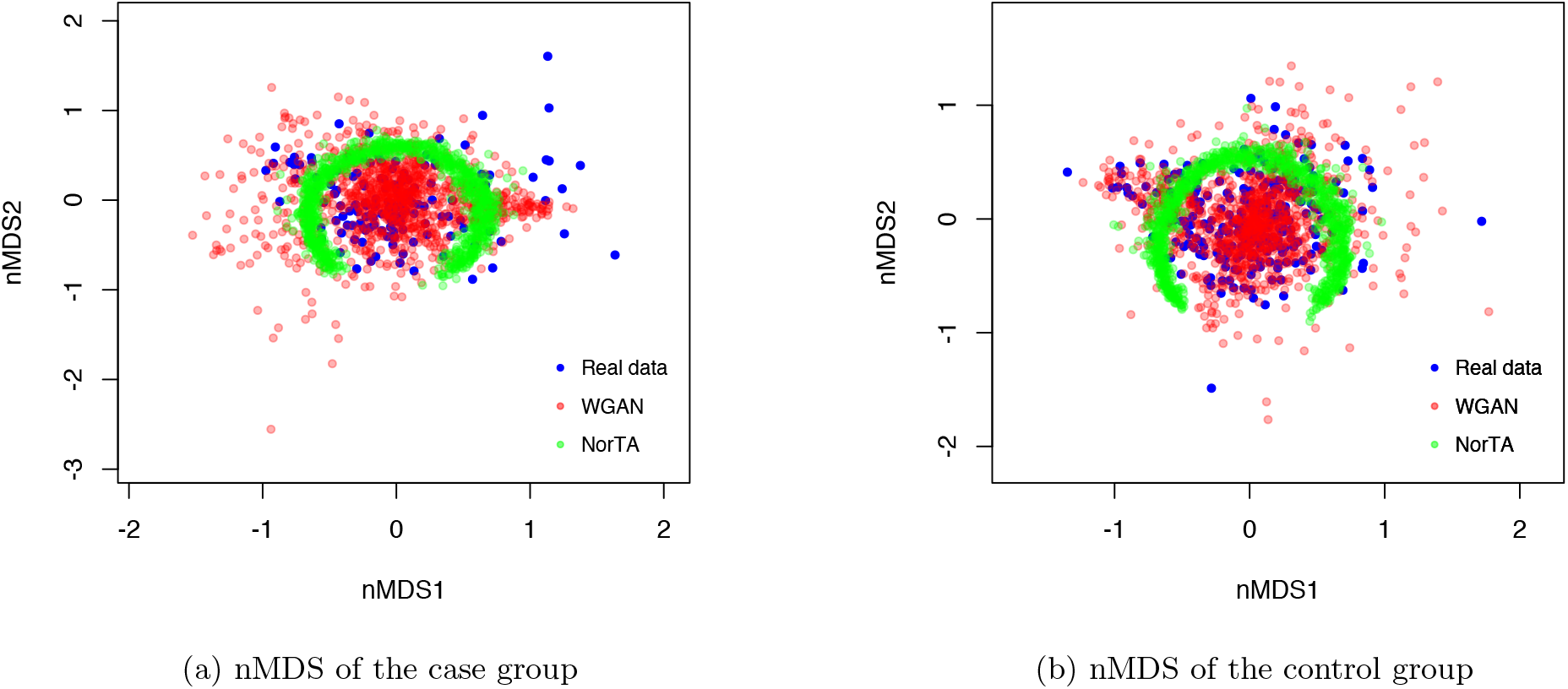
*β*-diversity visualization using non-metric multidimensional scaling (nMDS) for (a) the case and (b) the control groups. For samples from the real data, the MB-GAN, or the Normal To Anything (NorTA) simulated data, the two-dimensional nMDS values were calculated using the Bray-Curtis metric

The correlograms in Figure 5 (a) present three correlation matrices. A blue ellipse represents a negative correlation, while a red one suggests a positive correlation. The darker the color, or the shorter the ellipse’s minor axis, the stronger the correlation between the corresponding taxa pair. In Figure 5 (a), the overall scale of the correlation coefficients from MB-GAN simulated data is slightly smaller than those calculated from the real data. However, the pattern observed in the MBGAN simulated data shows a clear consistency with the true correlation structure. Furthermore, MB-GAN is able to preserve the strongest correlation signals in the real data. NorTA, on the other hand, shows even weaker signals of the correlation coefficients, and an obvious disparity between its correlation structure and the true one. The above conclusions also hold for the control group, as similar results are shown in Figure 5 (b).

Next, we compare the empirical distributions of the correlation coefficients. Each subfigure in Figure 6 (a) and (b) overlays two histograms of the correlation coefficients, with the blue one from the real data, the red one from MB-GAN, and the green one from NorTA. Here, MB-GAN shows a clear advantage over the NorTA in two aspects: first, its overall shape better matches the true coefficients’ distribution; second, as discussed previously, its overall range of the correlation coefficients is more comparable with the range given by the true coefficients, while the species generated by NorTA show an overall weaker correlations. Based on these results, we conclude that MB-GAN performs better in capturing the second order characteristics, the correlation structure, observed in the real microbiome data.

**Figure 5:**
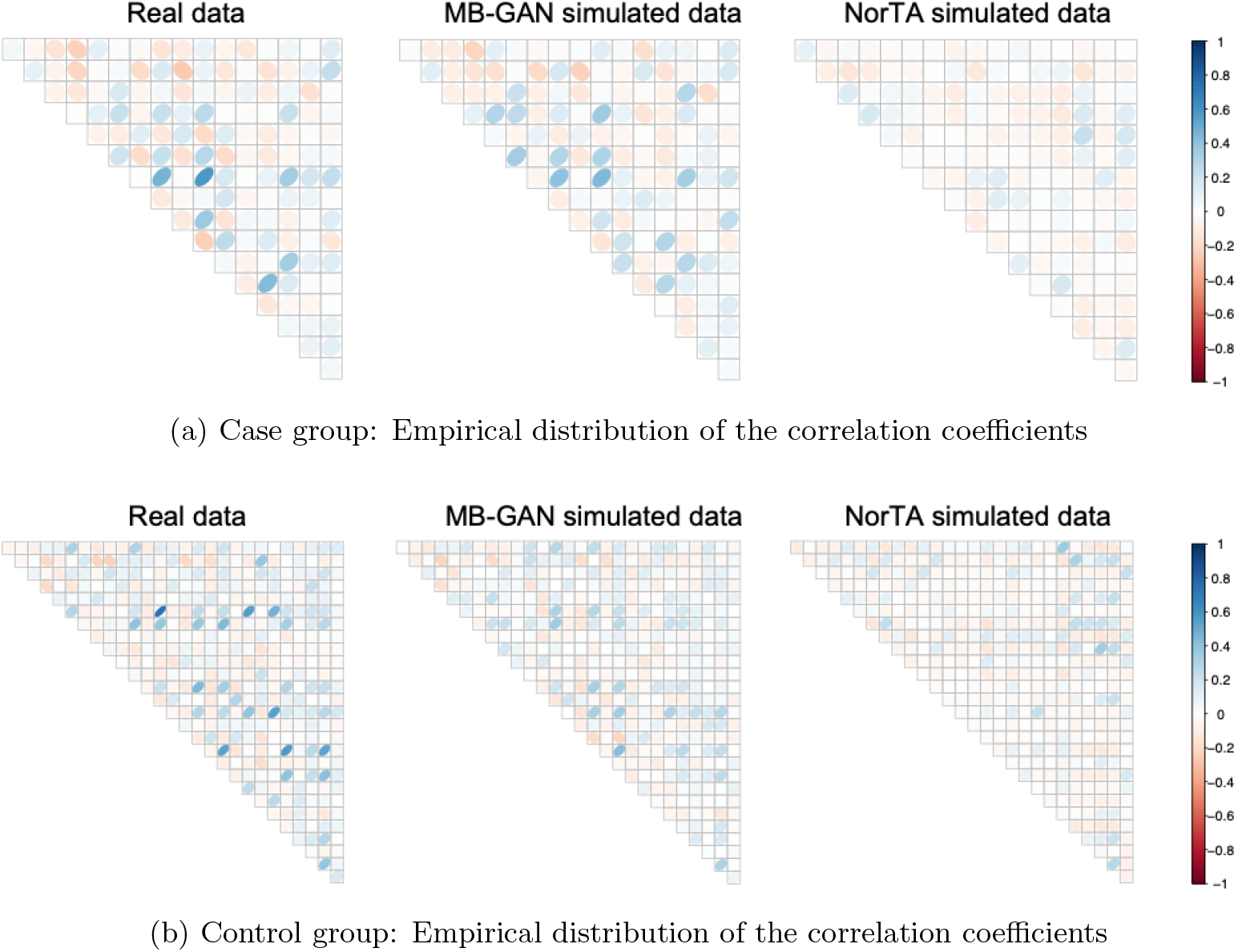
Comparison between the correlation matrices of the real data of the case group, the data simulated by MB-GAN and data simulated by Normal To Anything (NorTA). Results calculated from the case and the control groups are shown in (a) and (b), respectively.

### 3.4 Online resource for MB-GAN

We used Keras^[28]^ with the Tensorflow^[29]^ backend to implement the MB-GAN model. We provided the source codes, example output datasets and a Jupyter Notebook in GitHub as an online resource (https://github.com/zhanxw/MB-GAN). These are the trained models for the generator network and they can facilitate reproducing the results reported in this manuscript. They can also be customized to simulate new datasets for future microbiome studies. The codes are licensed under GNU General Public License (GPL) v3.0.

## 4 Discussion

We have developed a novel simulation framework, MB-GAN, for microbiome abundance data. It is designed based on the latest research into GANs. It can simulate microbiome data that are not easily distinguishable from the real data in terms of sample-level characteristics (sparsity, *α*-diversity and *β*-diversity) and taxa-taxa relationships. Furthermore, the simulator is trained by real data and does not need explicit statistical models.

The simulated data from the MB-GAN framework can facilitate microbiome research. For example, researchers can assign different effect sizes for the simulated taxa and develop differential abundance analysis methods used in MWAS where a group of taxa can be jointly tested ^[30]^. As MB-GAN can preserve taxa-taxa relationships, the simulated data can also facilitate microbiome network studies or microbial community studies. However, MB-GAN should not be used as a tool to enlarge existing sample sizes to detect differentially abundant taxa. Instead, it is preferred to perform MWAS after imposing disease models on the simulated case and control samples.

**Figure 6:**
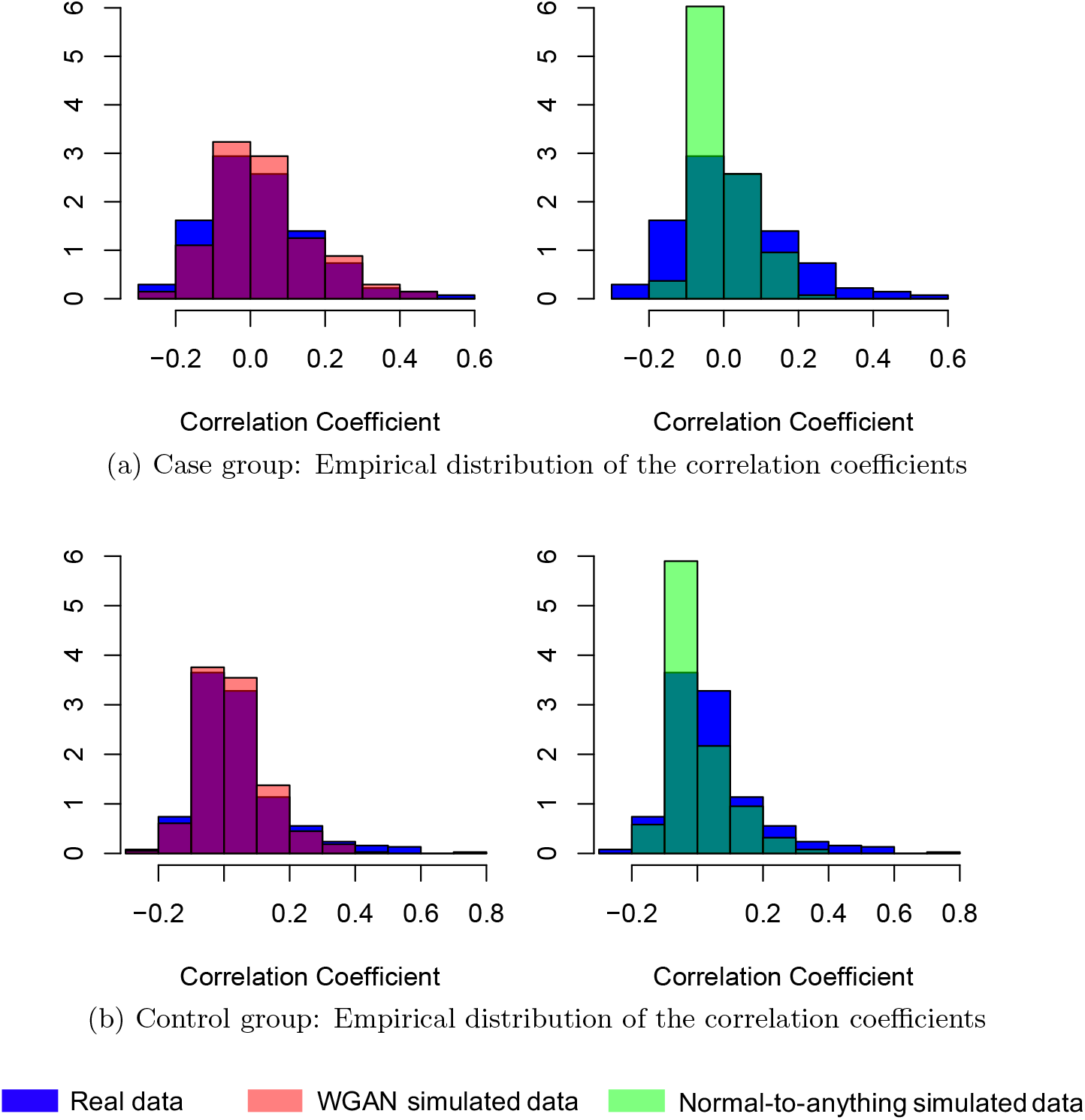
Comparison between the empirical distributions of the Pearson correlation coefficients calculated from the real data and the two types of simulated data of (a) case group, and (b) control group. The correlation coefficients are calculated from the species with less than 10% zeros across all samples in the corresponding group.

## 5 Acknowledgements

We thank Jessie Norris for her comments on the manuscript.

## 8 Author Contributions

R.R and S.J performed the experimental, L.X, G.X, Y.X, D.J.L. and Q.L. provided resources and helpful discussions, R.R., S.J. and X.Z. designed the experiment, performed data analysis, wrote software and wrote the manuscript.

